# Effects of Combined Viral and Bacterial Vaccine on Production Performance in Broiler Chickens

**DOI:** 10.64898/2025.12.27.696645

**Authors:** Qiang Han, Nan Wang, Teng Feng, Yue-Yue Li, Zhuang Zhu, Teng-Lin Xu, Ke-Ji Quan, George Fei Zhang

## Abstract

This study formulated a novel inactivated vaccine containing Newcastle disease virus (Lasota strain), avian influenza virus (BZ strains), APG, adenovirus and ORT. The vaccine was administered to 1-day-old and 7-day-old fast-growing broiler chickens. Experimental results showed that the vaccine reduced the feed conversion ratio and induced high levels of antibodies against Newcastle disease (Lasota strain), avian influenza (BZ strains), and avian rhinotracheitis. Immunization at 1 day of age had a significant impact on the production of antibodies against adenovirus. Challenge tests indicated that the vaccine partially protected the broiler chickens against Avibacterium paragallinarum types A, B, and C in broilers. Post-mortem examination of the injection sites showed good absorption with no significant unabsorbed oil droplets or hyperplasia in both 1-day-old and 7-day-old immunized chickens. This study demonstrated that combined viral and bacterial vaccine enhanced the feed conversion ratio in broiler chickens.

## 1 Introduction

Literature reports indicate that infection in broiler chickens leads to slow weight gain, causing significant economic losses. *Ornithobacterium rhinotracheale* (ORT) can co-infect broiler chickens of different ages, exacerbating disease symptoms. Therefore, this experiment was designed to investigate the effects of adding APG and ORT on broiler chicken weight, feed conversion ratio, and Newcastle disease antibody levels. Infectious pathogens could be identified by microbiota analysis (1, 2). Vaccination can protect the host from infectious disease and improve the production performance (3-6). In this study, we demonstrated the vaccination of this viral and bacterial combined vaccine in boiler chicken.

## 2 Materials and Methods

### 2.1 Bacteria and animals

APG-A, APG-B, APG-C are stored in the R&D Incubation Center’s bacterial and viral strain bank. Experimental animals: Healthy SPF chickens (Merial) older than 42 days old, and healthy conventional chickens older than 40 days old, provided by the R&D Incubation Center’s Wohua Bioeng Experimental Animal Center. Newcastle disease and adenovirus antibody levels: Tested by the testing center. Newcastle disease antibody levels were determined by ELISA method, and adenovirus antibody levels by agar gel immunodiffusion method. ORT antibody levels: ELISA method. Mineral oil, phosphate-buffered saline (PBS), 0.5% coenzyme solution (G/V, Roche), chicken serum (Kangyuan Biotechnology), TSA culture medium (BD), semi-synthetic culture medium (casein peptone, multi-valent peptone from Obio, yeast extract from Oxoid, monosodium glutamate from Taile, sodium chloride from Sinopharm, dipotassium hydrogen phosphate trihydrate from Tianjin Tianli), etc.

### 2.2 Vaccination

Viruses and bacteria were prepared for vaccine. Isolation chamber. Newcastle disease virus, avian influenza virus, adenovirus, APG, and ORT, Two hundred one-day-old fast-growing broiler chickens (AA broilers) were randomly divided into four groups. The initial weight of each group was recorded before immunization. Group 1 was immunized subcutaneously in the neck with 0.5 mL of Newcastle disease and infectious bronchitis vaccine at 1 day of age; Group 2 was immunized subcutaneously in the neck with 0.5 mL of Newcastle disease and infectious bronchitis vaccine at 7 days of age; Group 3 was immunized subcutaneously in the neck with 0.5 mL of Newcastle disease vaccine at 7 days of age; and Group 4 served as a control group. After immunization, the chickens’ mental state, posture, feeding, and drinking behavior were observed every two hours until the mental state of the control group and the immunized groups were consistent.

### 2.3 Weight gain measurement and Meat-to-fat ratio determination

After immunization, each group was fed a weighed amount of feed. At the end of the experiment, the feed consumption of each group was recorded, and the feed conversion ratio was calculated. The weight of the chickens in each group was recorded at the beginning of the experiment. The weight of each group was recorded every week, and the weight gain was calculated. Before the end of the experiment, the weight of the chickens in each group was weighed to compare whether the vaccine had an effect on chicken weight gain.

### 2.4 Antibody level testing

Before the end of the experiment, blood samples were collected from each chicken in each group. Agar gel immunodiffusion was used to determine the antibody levels against infectious coryza types A, B, and C. Indirect ELISA was used to detect antibody levels against the ORT, HI assay was used to detect Newcastle disease antibody levels, and agar gel immunodiffusion was used to detect adenovirus antibody levels.

### 2.5 Challenge assay

Before the end of the experiment, each group randomly selected eight chickens to be challenged with type A, B, and C infectious coryza to test the protective efficacy against the disease.

### 2.6 Statistical analysis

Graphpad was used to process the data.

## 3 Results

### 3.1 Humoral immune response detection

According to the Newcastle disease antibody level test results (Figure 2), the XLBXN-1d vaccine group affected Newcastle disease antibody levels, while there was no significant difference between the XLBXN-7d vaccine group and the XLX-7d group. The addition of APG and ORT affected the production of Newcastle disease antibodies. According to the avian influenza antibody test results (Figure 3), the XLBXN-1d vaccine group affected Newcastle disease antibody levels, while there was no significant difference between the XLBXN-7d vaccine group and the XLX-7d group. Immunization on day 1 affected avian influenza antibody production. According to the adenovirus agar gel immunodiffusion antibody test results (Figure 4), the XLBXN-1d vaccine group affected adenovirus antibody levels, the XLBXN-7d vaccine group performed better than the XLBXN-1d vaccine group, but the difference from the XLX-7d group was still significant. Immunization with the XLBXN vaccine on days 1 and 7 affected adenovirus antibody levels. The XLBXN-1d and XLBXN-7d vaccine groups both showed positive results for APG A, B, and C antibodies, indicating no effect on rhinotracheitis antibody production. The XLBXN-1d and XLBXN-7d vaccine groups both showed positive results for ORT antibodies, indicating no effect on ORT antibody production. However, the antibody levels in the day 7 immunization group were significantly higher than those in the day 1 immunization group.

**Figure 1:**
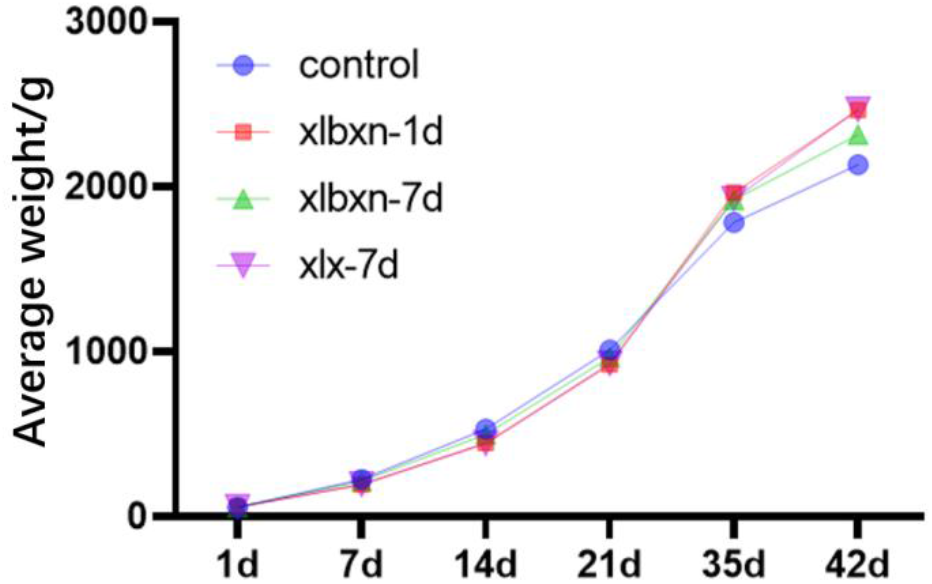
Average weight gain results

**Figure 2:**
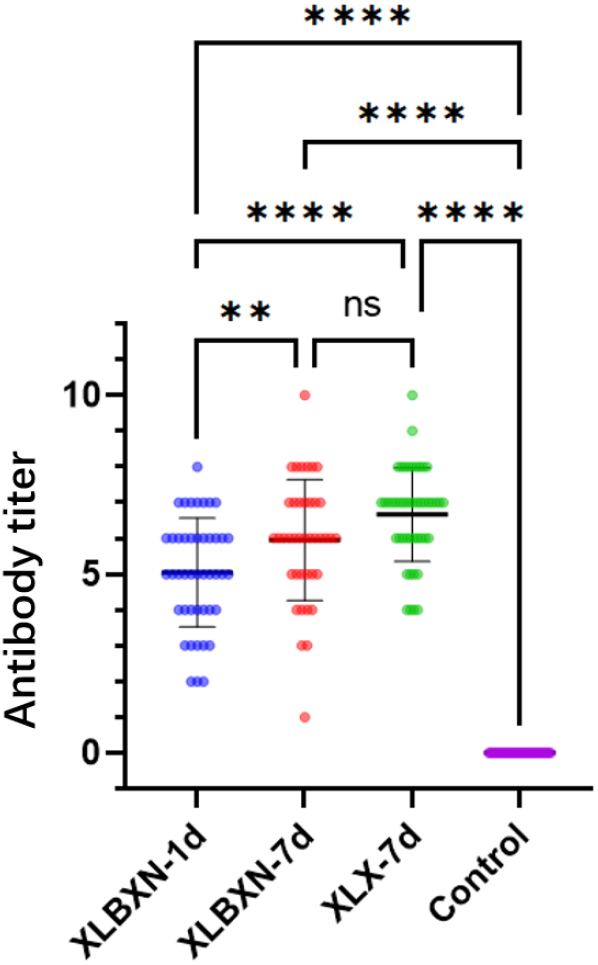
Newcastle disease antibody levels

**Figure 3:**
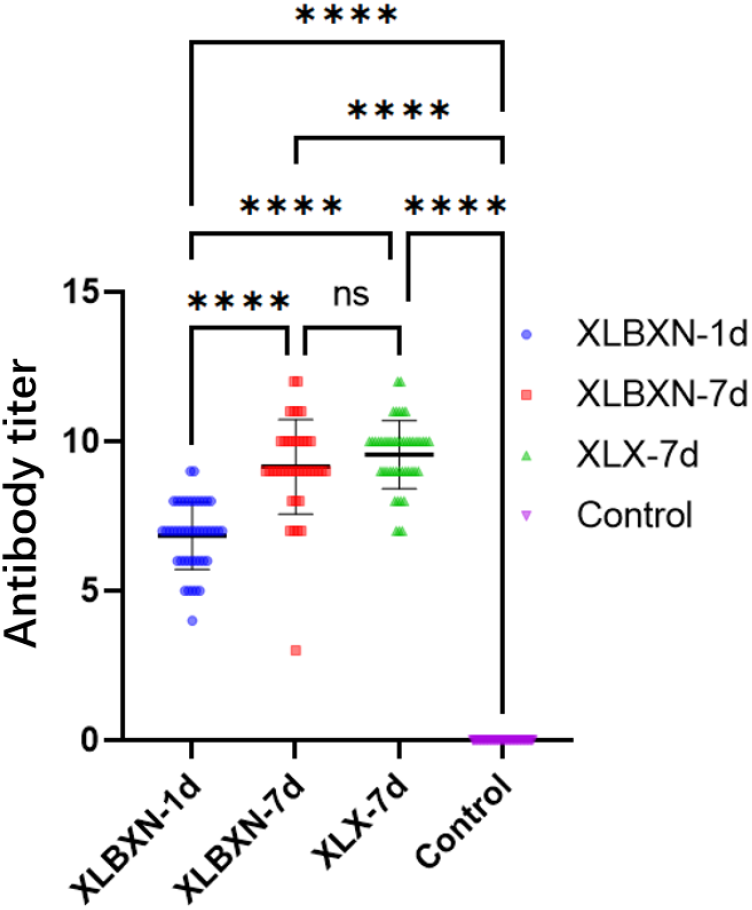
Avian influenza antibodies

**Figure 4:**
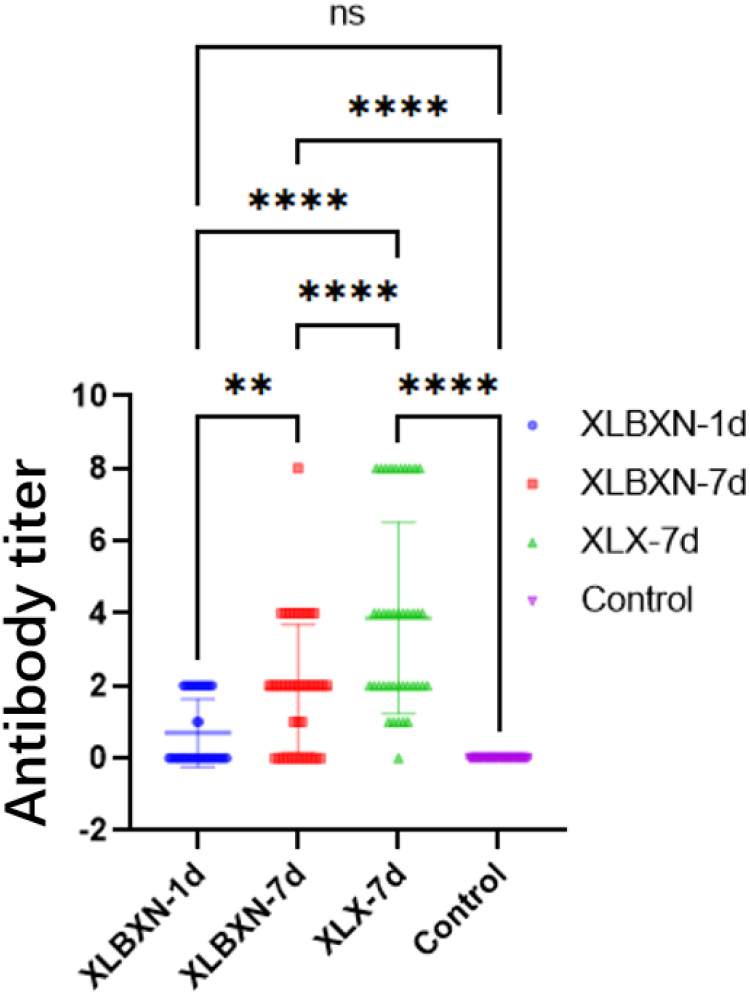
Adenovirus antibody

**Figure 5:**
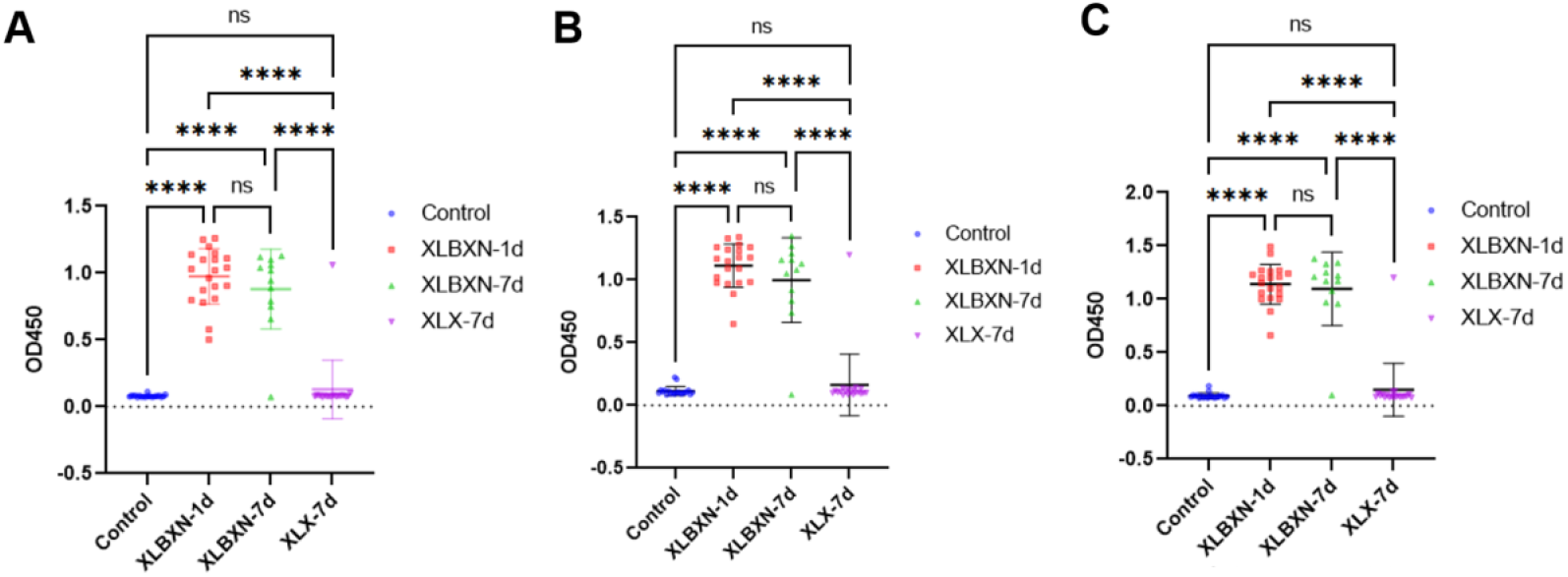
Infectious coryza ELISA antibody

**Figure 6:**
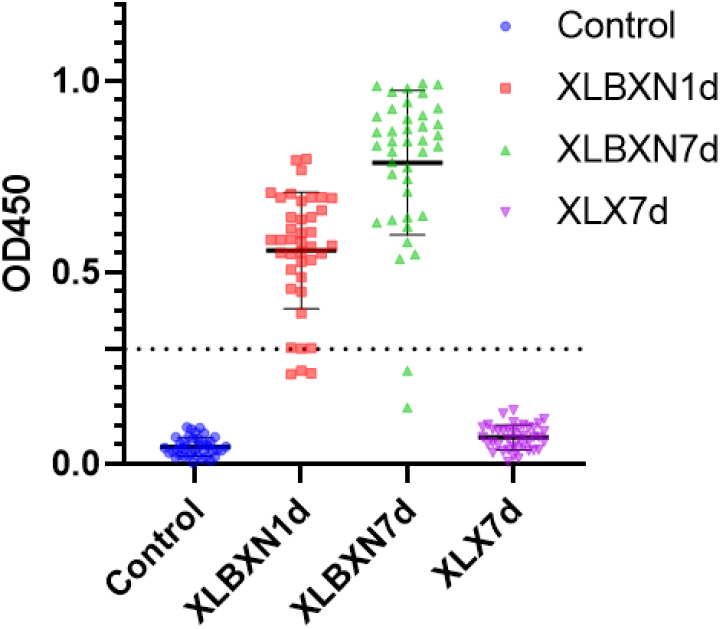
ORT antibody test

### 3.2 Weight gain measurement

The average body weight of broilers is shown in Figure 1. The control group was in the leading position at 14 and 21 days post-immunization, with significantly higher body weight than the other three immunized groups. At 35 days, the average body weight of the immunized groups exceeded that of the control group, and at 42 days of age, the XLBXN group immunized on day 1 and the XLX group immunized on day 7 were comparable and higher than the XLBXN group immunized on day 7. The average weight gain and feed intake of each group are shown in Table 1. Based on the feed conversion ratio (kg/kg) = feed intake/weight gain, the results are shown in Table 1. The feed conversion ratio of XLBXN-1d and XLBXN-7d was 0.1 lower than the control group, and XLX-7d was 0.03 lower than the control group. Overall, the feed conversion ratio of the immunized groups was lower than that of the control group.

**Table 1:**

|  | Control | XLBXN-1d | XLBXN-7d | XLX-7d |
| --- | --- | --- | --- | --- |
| Total weight (kg) | 104.5563 | 123.1777 | 115.829 | 120.9397 |
| Feed(kg) | 170.05 | 188.2 | 177.7 | 193.7 |
| Feed conversion ratio (kg/kg) | 1.63 | 1.53 | 1.53 | 1.6 |

### 3.3 Results of protection against rhinitis-causing pathogens

According to the challenge test results for APG types A, B, and C (Figure 7), the XLBXN-1d vaccine group showed good uniformity, with 4 animals protected against each serotype. The XLBXN-7d vaccine group showed better protection against type A (8 animals protected), but poorer protection against types B and C.

**Figure 7:**
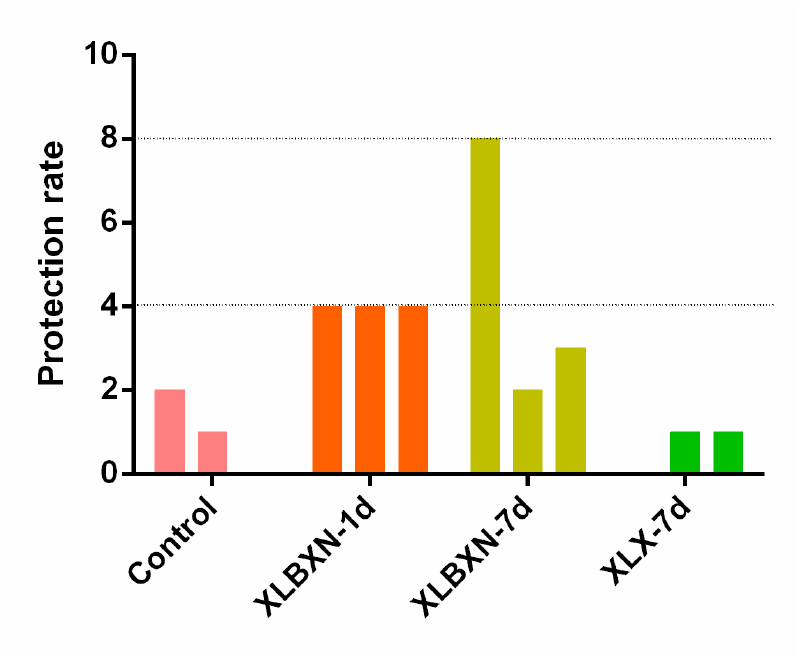
Challenge protection experiment for infectious coryza in chickens

## 4 Discussion

The addition of APG-A, APG-B, APG-C, and ORT to the feed reduced the feed conversion ratio of broilers by 0.1 percentage points. This reduction can significantly decrease feed costs in broiler farming. This reduction may be due to the increased disease resistance of broilers during growth as a result of vaccination, leading to a stronger immune system; or the vaccination may have altered the intestinal microflora of the broilers, thereby enhancing nutrient absorption.

However, areas for improvement are also evident. The addition of the five proteins affected the antibody levels against Newcastle disease, possibly due to the high protein content. Further optimization of the protein dosage is needed to mitigate the impact on Newcastle disease antibody levels. The inactivated avian influenza and Newcastle disease vaccine demonstrated good safety in broilers, and immunization at 7 days of age resulted in high antibody levels against various pathogens.

In conclusion, the combined viral and bacterial vaccine is safe for broiler chickens. This study demonstrated that combined viral and bacterial vaccine enhanced the feed conversion ratio in broiler chickens.

## Acknowledgment

This study was funded by the Taishan Industrial Expert Programme of Shandong Province (NO. tscx202306107). This study was supported by the Sichuan Natural Science Foundation (2025ZNSFSC1084).

## Author’s contribution

GF Zhang conceived and designed the study. Q Han, N Wang, T Feng, and YY Li executed the experiment. GF Zhang processed data analysis and writing. GF Zhang, KJ Quan, TL Xu, and Z Zhu edited the manuscript. All authors approved the final version of the manuscript.

## Declaration of conflicting interest

The author(s) declared no potential conflicts of interest with respect to the research, authorship, and/or publication of this article.

## Data availability statement

The raw data supporting the conclusions of this article are available from the authors upon reasonable request.

## Notes

### Competing Interest Statement

The authors have declared no competing interest.

## References

1. Han Q, Wang N, Feng T, Li Y-Y, Zhu Z, Xu T-L, Quan K-J, Zhang GF. 2025. Characteristics of reproductive tract microbiota in health and disease. bioRxiv doi:10.64898/2025.12.20.695662.

2. Li MN, Wang T, Wang N, Han Q, You XM, Zhang S, Zhang CC, Shi YQ, Qiao PZ, Man CL, Feng T, Li YY, Zhu Z, Quan KJ, Xu TL, Zhang GF. 2024. A detailed analysis of 16S rRNA gene sequencing and conventional PCR-based testing for the diagnosis of bacterial pathogens and discovery of novel bacteria. Antonie Van Leeuwenhoek 117:102.

3. Zhang F, Zhang Y, Wen X, Huang X, Wen Y, Wu R, Yan Q, Huang Y, Ma X, Zhao Q, Cao S. 2015. Identification of Actinobacillus pleuropneumoniae Genes Preferentially Expressed During Infection Using In Vivo-Induced Antigen Technology (IVIAT). J Microbiol Biotechnol 25:1606–13.

4. Zhang F, Zhao Q, Quan K, Zhu Z, Yang Y, Wen X, Chang YF, Huang X, Wu R, Wen Y, Yan Q, Huang Y, Ma X, Han X, Cao S. 2018. Galactose-1-phosphate uridyltransferase (GalT), an in vivo-induced antigen of Actinobacillus pleuropneumoniae serovar 5b strain L20, provided immunoprotection against serovar 1 strain MS71. PLoS One 13:e0198207.

5. Zhang GF, Meng W, Chen L, Ding L, Sun S, Wang X, Huang Y, Guo H, Gao SJ. 2023. Infectivity of pseudotyped SARS-CoV-2 variants of concern in different human cell types and inhibitory effects of recombinant spike protein and entry-related cellular factors. J Med Virol 95:e28437.

6. Zhang N, Quan K, Lin M, Lu Z, Li Z, Yang Y, Xu N, Yang H, Zhu J, Zhang GF. 2025. Comprehensive evaluation of HA epitope modifications in H9N2 subtype avian influenza vaccines for broad cross-protection. Journal of Integrative Agriculture.

